# The Cortical Output System that Controls a Single Vibrissa Muscle

**DOI:** 10.1101/2025.03.18.643882

**Authors:** Aman Maharjan, Jason M. Guest, Jean-Alban Rathelot, Fiorella M. Gomez Osorio, Peter L. Strick, Marcel Oberlaender

## Abstract

What is the neural substrate that enables the cerebral cortex to control a single mystacial vibrissa and orchestrate its movement? To answer this question, we injected rabies virus into the intrinsic muscle that protracts the rat C3 vibrissa and used retrograde transneuronal transport to identify the cortical neurons that control the muscle. A surprisingly diverse set of cortical areas is the origin of disynaptic control over the motoneurons that influence the C3 protractor. More than two thirds of these layer 5 pyramidal neurons (L5PNs) are dispersed in frontal and parietal areas outside the primary motor cortex (vM1). This observation emphasizes the importance of descending commands from non-primary motor areas. More than a third of the L5PNs originate from somatosensory areas, such as the barrel field (vS1). The barrel field has been long considered a prototypic model system for studying sensory processing at the level of the cerebral cortex. Even so, we find that the number of L5PNs in vS1, and even their peak density, rivals the number and peak density of L5PNs in vM1. Thus, our results emphasize the importance of the barrel field in processing motor output. The distribution of L5PNs in vM1 and vS1 leads us to propose a new model of vibrissa protraction in which vM1 output results in protraction, and vS1 output results in reciprocal inhibition (suppression) of protraction. This paired initiation and suppression of complementary movements may be a general feature of the descending control signals from the rodent M1 and S1.

## Introduction

The facial vibrissae are quintessential sensorimotor organs for rodents (1, 2). On the sensory side, tactile information from single vibrissae is processed by anatomically segregated clusters of neurons in the brainstem, thalamus and cerebral cortex (3). Indeed, layer 4 (L4) of the vibrissal-related part of the primary somatosensory cortex (vS1) contains a “barrel field” with an anatomically distinct cluster of neurons (i.e., “barrel”) for each vibrissa (4). The vibrissal system is widely used as a model for investigating the anatomical and physiological substrates of sensory processing at the cortical level (5).

The vibrissae motor side is equally interesting (6). There at least two modes of motor control for the vibrissae. During “whisking”, the vibrissae on both sides of the face are actively moved back and forth in concert to gather tactile information about the environment (2). In a second mode of motor control, single vibrissa are moved in highly complex, individuated patterns as part of such an active sensing process (2). These modes of motor control are generated largely by two classes of muscles. “Extrinsic” muscles evoke backward movement – retraction – of all the vibrissae within a single row (7). In contrast, “intrinsic” muscles evoke forward movement – protraction – of a single vibrissa (8), and thereby enable the complex, yet highly flexible, individual vibrissa movements during active sensing (9). Here we reveal the anatomical substrate that enables control of the protractor muscle of a single vibrissa. We focus our investigation on the cerebral cortex, because it is considered to be a key brain region that endows animals with the ability to precisely control muscles and produce complex movement patterns (10).

An early study suggested that some neurons in layer 5 (L5) of the primary motor cortex (M1) form direct monosynaptic connections with the motoneurons (MNs) of vibrissa muscles (11), but this has subsequently been ruled out (6, 10, 12). Instead, the cerebral cortex influences vibrissa MNs via disynaptic pathways that are mediated by distinct sets of “last-order” interneurons – i.e., premotor neurons (preMNs), located at multiple sites in the brainstem. These sites include several motor nuclei of the reticular formation, sensory nuclei of the spinal trigeminal complex, and nuclei of the respiratory and vestibular systems (12-19). The dispersed nature of these preMNs, and their complex patterns of inputs and outputs (12, 17), has made it especially challenging to dissect the disynaptic pathways that link the cerebral cortex to vibrissa MNs.

Here we overcome these challenges by using retrograde transneuronal transport of rabies virus (20) from a single vibrissa protractor muscle. Retrograde transport of the virus first infects the MNs (“1st-order” neurons) that innervate the injected muscle (12). Then, retrograde transneuronal transport infects all preMNs (“2nd-order” neurons) that innervate these MNs (12). By allowing another stage of transneuronal transport, rabies virus then infects all layer 5 pyramidal neurons (L5PNs, “3rd-order” neurons) that innervate these preMNs. Our results show that multiple cortical regions in both hemispheres, including the S1 barrel field, as well as M1, M2, S2 and the anterior insular cortex (AI) are the major sources of the disynaptic input from the cerebral cortex to the MNs of the protractor muscle of a single vibrissa.

## Results

We injected rabies virus into a single vibrissa muscle – i.e., the intrinsic muscle that protracts the C3 vibrissa of rats (n=8). The C3 vibrissa is located in the approximate center of the mystacial pad **(Fig. 1A)**. We varied the survival time between 3-5 days and then mapped the number and location of neurons infected with rabies virus in each of the brains **(Fig. S1A-E)**.

**Figure 1.**
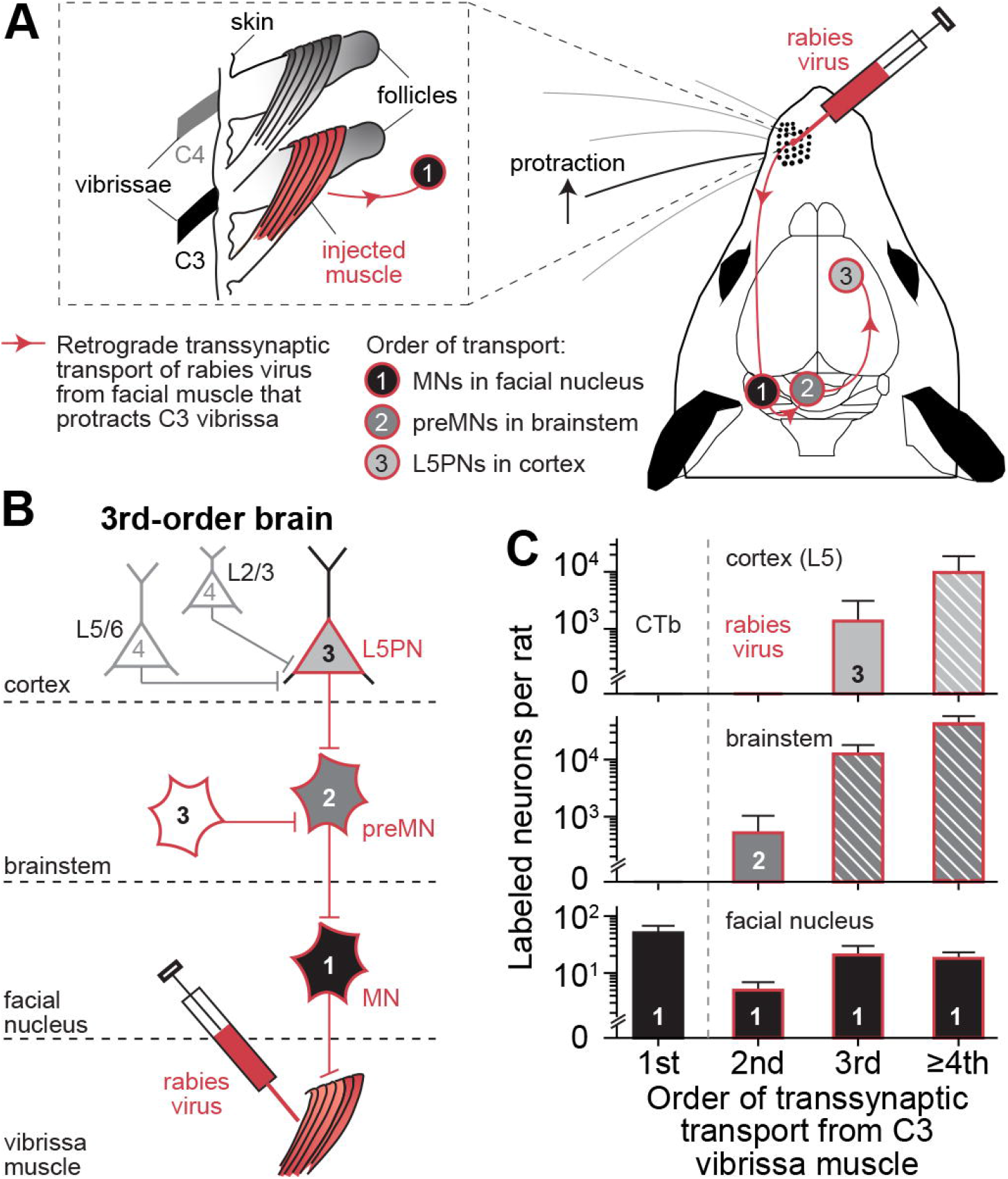
Schematic illustration of experimental approach. **A**. We injected the N2c strain of rabies virus into a single intrinsic muscle that protracts the C3 facial vibrissa of rats. **B**. Depending on the survival time, retrograde transsynaptic transport from the muscle infects 1st-order motoneurons (MNs) in the ventrolateral facial nucleus, 2nd-order premotor neurons (preMNs) in the brainstem, 3rd-prder pyramidal neurons in cortical layer 5 (L5PNs) and ≥4th-order neurons in other cortical layers (e.g. in L2/3). Thus, in brains where transsynaptic transport was limited to 3rd-order, cortical neurons infected with rabies virus are restricted entirely to L5. **C**. Numbers of retrogradely labeled neurons (mean ± STD) in the facial nucleus, brainstem and L5 of 1st- (n=10), 2nd- (n=2), 3rd- (n=3) and ≥4th-order brains (n=3) following injections into the C3 vibrissa muscle. 1-st order data represents C3 injections of Cholera Toxin subunit B (CTb) that we reported in part previously (12). Dashed boxes for brainstem represent mix of preMNs and other neurons infected with rabies virus at ≥3rd-order. Dashed box for cortex represents mix of L5PNs with disynaptic connections to vibrissa MNs and other L5 neurons infected with rabies virus at ≥4th-order.

MNs that drive intrinsic muscles are located in the ventral part of the lateral facial nucleus in the brainstem (21), where they are spatially segregated from the more dorsally located MNs of extrinsic muscles (19). Each intrinsic muscle is innervated by a unique set of MNs (12, 21). These MN pools for each intrinsic muscle are arranged in an orderly map along the medial-lateral axis of the facial nucleus (12, 21). Our prior (12) and current results using retrograde transport of Cholera Toxin subunit B (CTb) demonstrate the precise location and size of the MN pool of the intrinsic muscle that protracts the C3 vibrissa **(Fig. S1F)**. This pool contains, on average, 48 MNs (median/25th/75th percentile: 51/42/58; n=10). Retrograde transport of rabies virus infected, on average, 16 MNs that were restricted to the appropriate location of the C3 MN pool **(Fig. S1F)**. Thus, our experiments define the inputs to about 1/3rd of the MN pool that protracts the C3 vibrissa.

The number and location of infected MNs remained relatively constant across animals and survival times (median/25th/75th percentile of MNs: 16/8/23; n=8). This observation emphasizes two points. First, rabies virus did not spread beyond the injection site to infect the MNs of other muscles (12). Second, the MNs of the intrinsic muscle of the C3 vibrissa are not interconnected with each other or with other MN pools (12). These results provide further evidence that the neurons infected with rabies virus at all “orders of retrograde transsynaptic transport” (i.e., the number of synapses crossed) are specifically related to the control of MNs that protract the C3 vibrissa.

We determined the order of transsynaptic transport based on the brain-wide distribution of the neurons infected with rabies virus **(Fig. 1B)**. In 2nd-order brains (n=2), rabies virus infected 1st-order MNs and 2nd-order preMNs in the brainstem **(Fig. S1A-B)**. In 3rd-order brains (n=3), rabies virus infected 3rd-order L5PNs in the cerebral cortex **(Fig. S1C-D)**, as well as 1st-, 2nd- and 3rd-order neurons in the brainstem **(Fig. S1E)**. Additional neurons infected with rabies virus were located in the midbrain and cerebellum of 3rd-order brains **(Table S1/2)**. In ≥4th-order brains (n=3), rabies virus infected neurons not only in L5 of the cerebral cortex, but also in other cortical layers **(Fig. 1C)**. The focus of this report is limited to 3rd-order brains in which cortical neurons infected with rabies virus were entirely limited to L5 **(Fig. S2A-C)**. In essence, these infected cortical neurons are L5PNs with disynaptic connections to MNs that control the protractor muscle of the C3 vibrissa – in the following referred to as vL5PNs.

One of our major results is the finding that multiple distinct areas of the cerebral cortex, in both hemispheres, contain vL5PNs **(Fig. 2A-B, Fig. S2A-B, Table 1)**. These vL5PNs are located not only in the primary and secondary motor areas of the cerebral cortex (M1 and M2), but also in large portions of the primary and secondary somatosensory cortex (S1 and S2), as well as in the anterior insular cortex (AI). Indeed, almost 2/3rds of the vL5PNs that control the protractor muscle of the C3 vibrissa are located outside of contralateral M1.

**Table 1.** Quantification of the total cortical output to control a single vibrissa protractor muscle and a single forelimb muscle (EDC). Numbers are percentages of the total L5PNs in the cerebral cortex.

| single vibrissa protractor |  |  |  | extensor digitorum communis |  |  |  |  |  |
| --- | --- | --- | --- | --- | --- | --- | --- | --- | --- |
| vM1 | 29.1% | vM1 | 13.7% | } 60.6% motor | fM1 | 43.5% | fM1 | 0.6% | } 46.9% motor |
| vM2 | 11.0% | vM2 | 6.8% |  | fM2 | 2.8% | fM2 | 0.0% |  |
| 40.1% motor |  | 20.5% motor |  |  | 46.3% motor |  | 0.6% motor |  |  |
| vS1 | 18.0% | vS1 | 1.0% | } 35.3% sensory | fS1 | 49.5% | fS1 | 0.2% | } 53.1% sensory |
| vS2 | 1.7% | vS2 | 3.1% |  | fS2 | 0.7% | fS2 | 0.0% |  |
| S1<sub>other</sub> | 8.9% | S1<sub>other</sub> | 2.6% |  | S1<sub>other</sub> | 2.1% | S1<sub>other</sub> | 0.8% |  |
| 28.6% sensory |  | 6.7% sensory |  |  | 52.3% sensory |  | 0.8% sensory |  |  |
| AI | 1.6% | AI | 2.5% | } 4.1% insular | AI | 0.0% | AI | 0.0% | } 0.0% insular |
| 1.6% insular |  | 2.5% insular |  |  | 0.0% insular |  | 0.0% insular |  |  |
| 70.3% contra |  | 29.7% ipsi |  |  | 98.6% contra |  | 1.4% ipsi |  |  |

**Figure 2.**
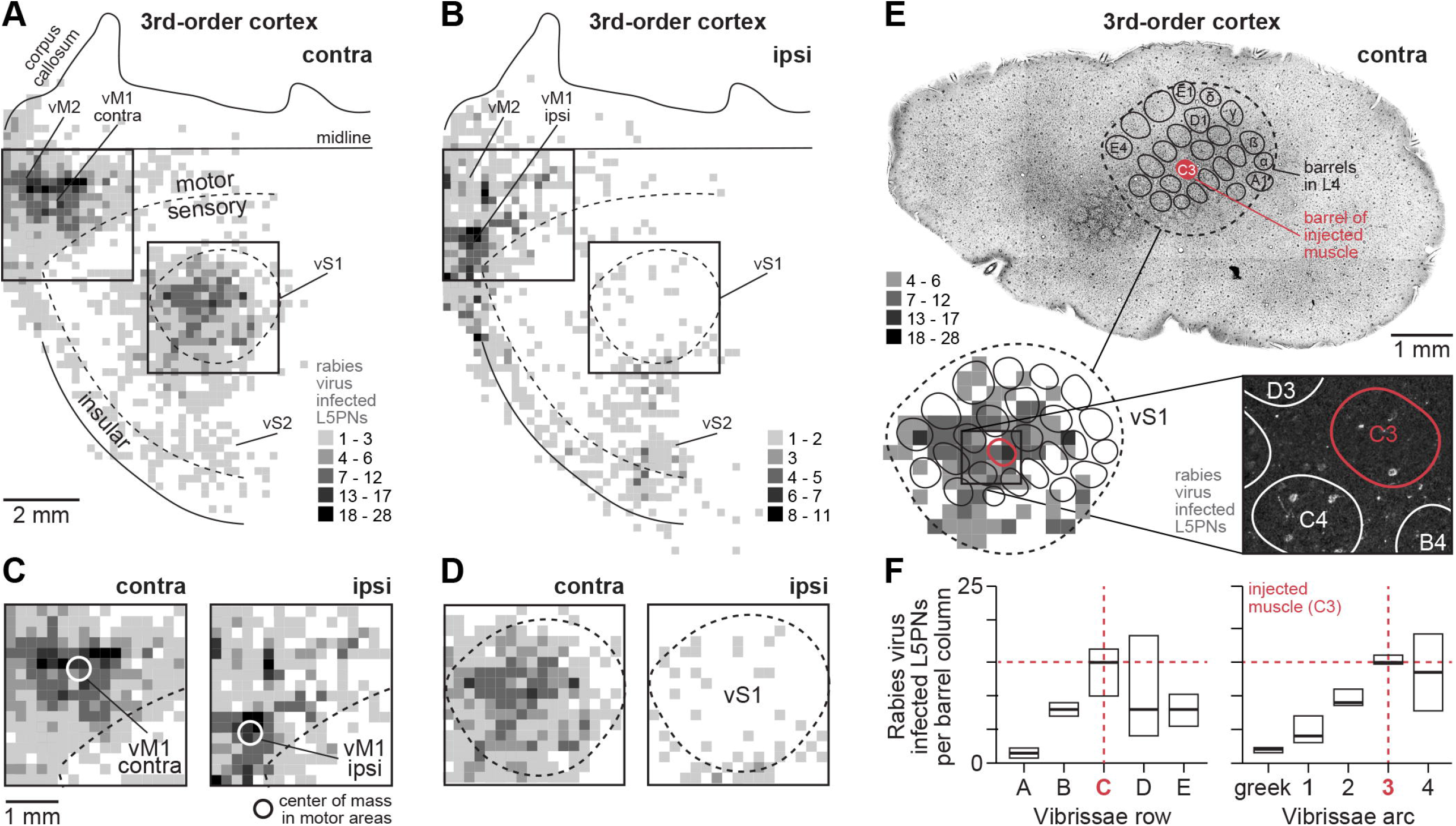
Cortical output system that controls a single vibrissa protractor muscle. **A**. Distribution of L5PNs infected with rabies virus (3rd-order, n=3). Data is visualized as L5PN density per 200µm x 200µm of a flat map of contralateral cortex (left-to-right: anterior-to-posterior). The corresponding 3D whole-brain reconstructions are shown in **(Fig. S2)**. Dashed lines represent cytoarchitectonic borders between motor, somatosensory and insular cortex **(Fig. S1C)**, and vibrissae-related primary somatosensory cortex (vS1, barrel cortex). **B**. Same as in panel A for ipsilateral cortex. **C**. Enlargement of boxes around motor areas in panel A (left) and panel B (right). White circles denote center locations of highest densities in contra- and ipsilateral primary motor cortex (vM1). **D**. Same as in panel C, but for boxes around vS1. **E**. Top: Brightfield image of 50µm thick section cut tangentially to vS1 for one of the brains from panel A-B. This section reveals the barrel field in L4. Barrel are outlined in black and labeled. Lower-left: Enlargement of the barrel field shows density from panel A (without lowest density bin). Bottom-right: Confocal image for a deeper section through L5 shows examples of rabies virus infected L5PNs with respect to barrel outlines (white). **F**. L5PNs per cortical barrel column (n=21, median ± 25th/75th percentile) for the brain in panel E. Barrel columns were grouped by vibrissae row (left) and arc (right).

A sizeable number of the vL5PNs (nearly 30%) are located in the hemisphere of the cerebral cortex that is ipsilateral to the injected vibrissa muscle **(Table 1)**. More than 2/3rds of these ipsilateral projecting vL5PNs are located in the cortical motor areas. The ratio of contralateral to ipsilateral projecting vL5PNs is ∼2 to 1 in both M1 and M2. These results may provide an anatomical basis for the bilateral nature of the cortical control over vibrissa protraction that is evoked by functional stimulation of the cortical motor areas (6, 22, 23).

The vL5PNs that are located in M1 are dispersed throughout the entire body map in each hemisphere **(Fig. 2A-B)**. In fact, small numbers of vL5PNs are located even in the hindlimb representations of contralateral and ipsilateral M1. Even so, the peak density of vL5PNs is located in the region of contralateral M1 that is commonly associated with vibrissa protraction (3). Surprisingly, another distinct peak density of vL5PNs is located in ipsilateral M1. We term these two dense regions vM1_contra_ and vM1_ispi_, respectively **(Fig. 2C)**. An overlap of the contralateral and ipsilateral maps of vL5PNs demonstrates that vM1_ipsi_ is located ∼1.5 mm more anterior and lateral than vM1_contra_ **(Fig. 3)**. The ratios of contralateral to ipsilateral projecting vL5PNs for vM1_contra_ is 15 to1, and the corresponding ratio for vM1_ipsi_ is 1 to 5. Taken together, these results have three important implications. First, the anatomical substrate exists for a broad region of M1 to influence the C3 vibrissa. On the other hand, M1 in each hemisphere contains separate hotspots for evoking protraction of the C3 vibrissa on both sides of the face. Finally, one hotspot, vM1_contra_, is biased towards protraction of the contralateral vibrissa, whereas the other relatively weaker hotspot, vM1_ipsi_, is biased towards protraction of the ipsilateral vibrissa.

**Figure 3.**
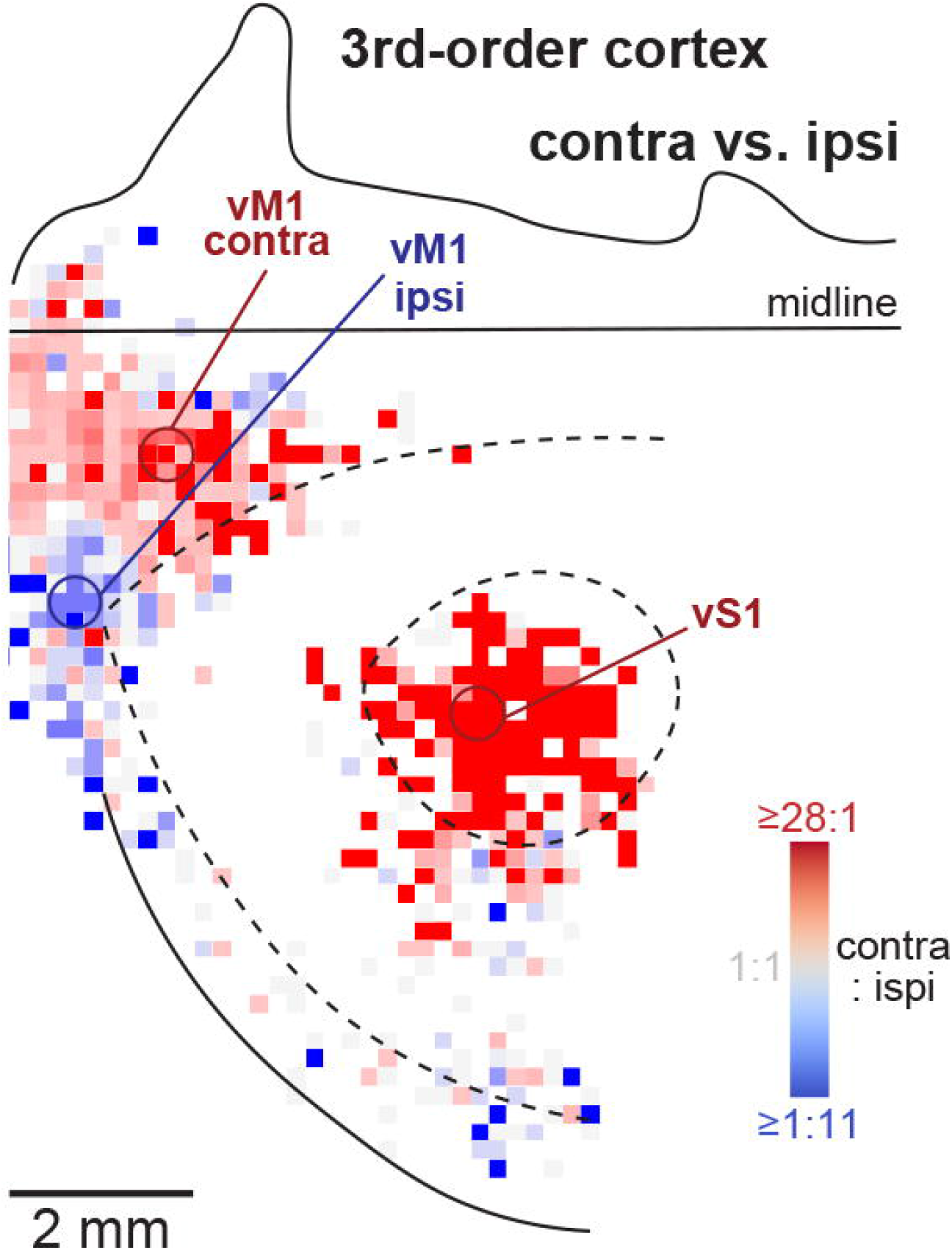
Contralateral vs. ipsilateral control of a single vibrissa protractor muscle. We generated this map by overlapping the contra- and ipsilateral density maps from **Fig. 2A/B**. The colors reflect the ratio between the L5PN density maps (without lowest density bins). Red colors represent overlap bins with higher numbers of L5PNs on the contralateral side than on the ipsilateral side. Blue colors represent overlap bins with higher numbers of L5PNs on the ipsilateral side than on the contralateral side.

Another major result of this study is that a sizeable fraction of the vL5PNs (just under 40%) is located in the somatosensory areas of the cerebral cortex **(Fig. 2A-B**). More than 50% of these somatosensory-related vL5PNs are located inside the barrel field (i.e., vS1) of the contralateral hemisphere **(Fig. 2D)**. Indeed, the barrel field stands out **(Fig. S3A)** as the sole cortical region with a nearly exclusive focus on the control of contralateral vibrissae – i.e., the ratio of contralateral to ipsilateral projecting vL5PNs is 80 to 1 in vS1. Our results emphasize two important features of vS1. First, it is a cortical “motor” area with a descending output capable of influencing the MNs of the C3 protractor muscle as directly as the output from M1. Second, it is the main cortical source of unilateral output to the protractor muscles.

We compared the cortical map of motor output to the MNs of a single vibrissa muscle with the cortical map of sensory input from single vibrissae. For this purpose, we reconstructed the barrel field of one of the 3rd-order animals **(Fig. 2E)**. This comparison shows that large numbers of vL5PNs are found not only below the barrel of the C3 vibrissa, but vL5PNs also extend below all barrels of the C-row, and the entire 3rd arc, as well as below barrels of the 2nd and 4th arcs in the B-, D- and E-rows **(Fig. 2F)**. This arrangement indicates that the motor output to influence protraction of the C3 vibrissa originates from approximately 70% of the barrel field. Thus, in contrast to the well-established one-to-one relationship between a single vibrissa and a single barrel on the sensory input side, the motor output side to a single vibrissa muscle is not one-to-one, but instead should be considered as “barrel field-to-one”.

In another set of rats (n=3), we injected rabies virus into a forelimb muscle **(Fig. 4A)**, the extensor digitorum communis (EDC). EDC functions to extend and spread the digits. We set the survival time to infect 3rd-order neurons. Consequently, cortical neurons infected with rabies virus were entirely limited to L5PNs **(Fig. S4A)**. These L5PNs have disynaptic connections to the MNs of the injected forelimb muscle (i.e., fL5PNs).

**Figure 4.**
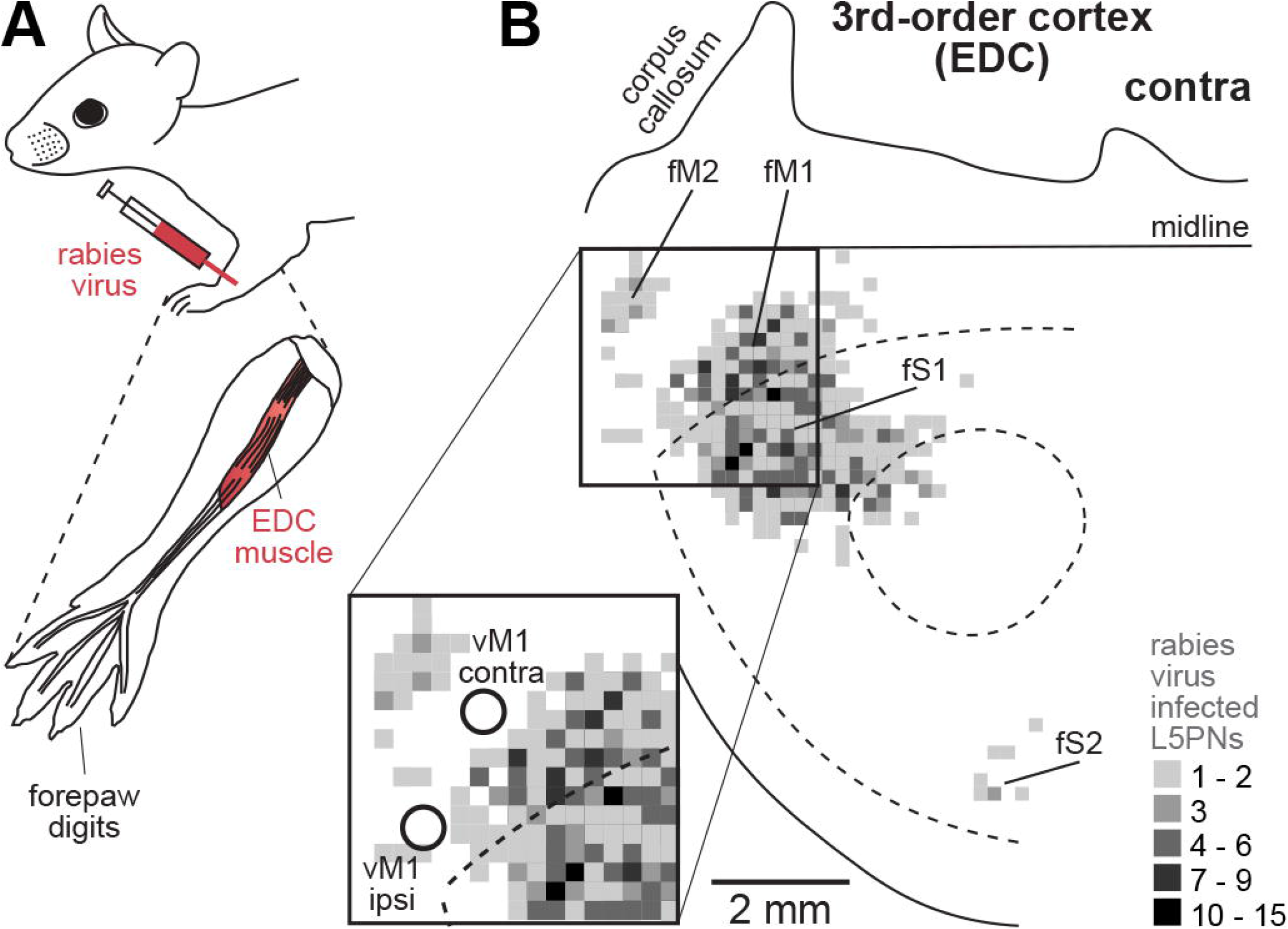
Cortical output system that controls a single forelimb muscle. **A**. We injected rabies virus into a single forelimb muscle that extends and spreads the digits, the extensor digitorum communis (EDC). **B**. Distribution of L5PNs infected with rabies virus (3rd-order, n=3). Map and borders as in **Fig. 2**. Enlargement shows the location of fM1 with respect to vM1 in contra- and ipsilateral cortex.

The distribution of fL5PNs differs from that of the vL5PNs in three striking ways **(Fig. 4B)**. First, none of the fL5PNs are located outside of the motor and somatosensory areas – i.e., none in insular cortex. Second, fL5PNs originate almost exclusively from the contralateral hemisphere **(Fig. S4B)** – i.e., the overall ratio of contralateral to ipsilateral projecting fL5PNs is 70 to 1 as compared to the overall ratio of 2 to 1 for vL5PNs. Third, forelimb S1 (fS1) contains a relatively greater proportion of the L5PN population than vibrissa S1 (i.e., 50% of the fL5PNs are located in fS1, 18% of the vL5PNs are located in vS1) **(Table 1)**. These results imply dramatic differences in the organization of the cortical control of vibrissae and forelimb muscles.

To examine potential interactions between cortical output systems for vibrissa and forelimb motor control, we overlapped the contralateral maps of vL5PNs and fL5PNs **(Fig. 5A)**. This overlap revealed remarkable intermingling of the two systems in M1 and M2. In fact, 84% of the fL5PNs are found in vM1. 38% of the vL5PNs are found in fM1. In M2, fL5PNs intermingle entirely within the more extensive group of vL5PNs in M2. In contrast, intermingling in S1 is much less substantial, where overlap occurs only at the vS1 to fS1 border. Thus, the somatosensory and motor areas of the rat cerebral cortex have different rules for vibrissa and forelimb representation: extensive overlap in motor areas vs. distinct representations in somatosensory areas **(Fig. 5B)**.

**Figure 5.**
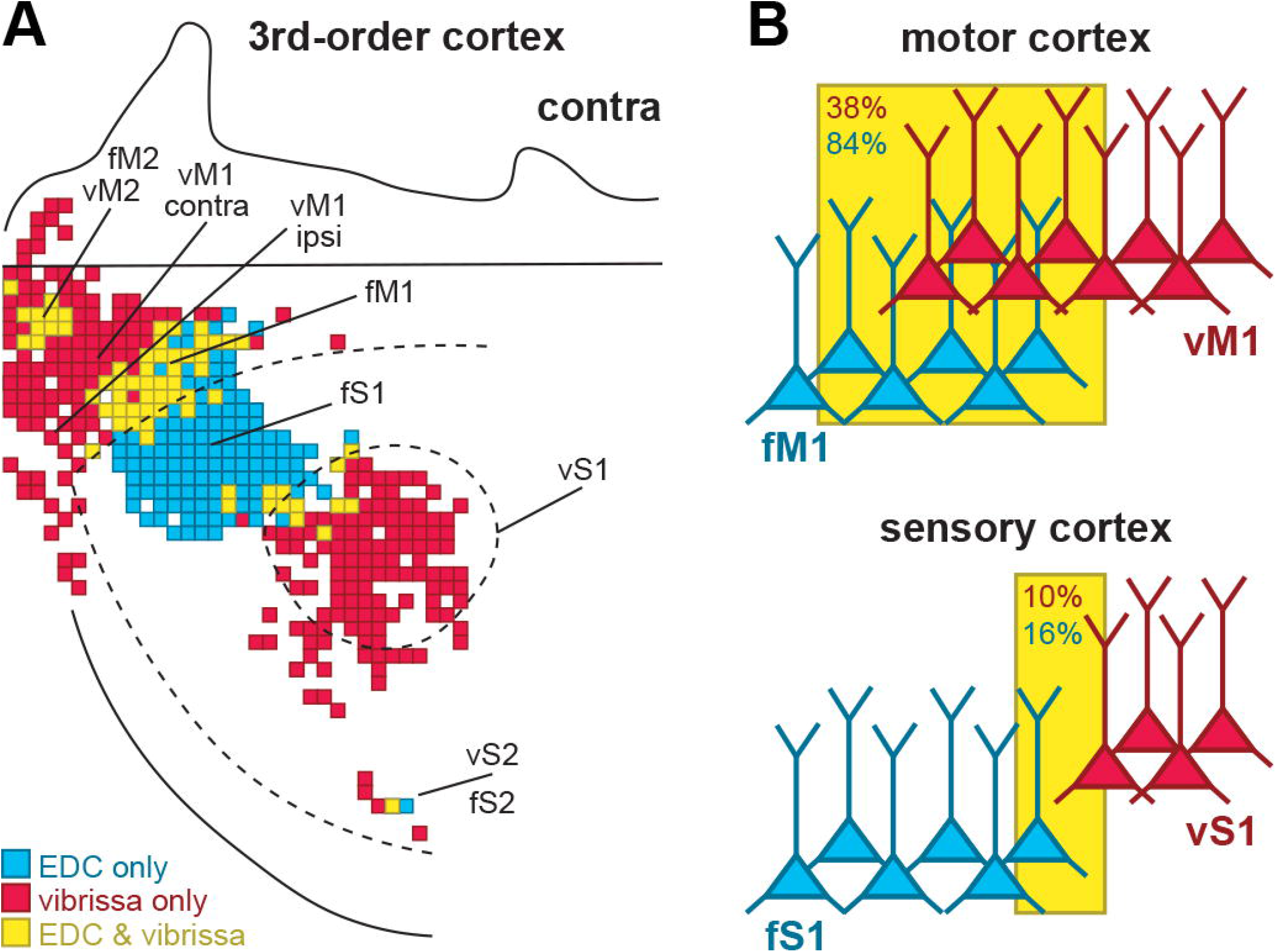
Cortical output system of a single vibrissa vs. a single forelimb muscle. **A**. Overlap of the L5PN density maps of the contralateral hemisphere from **Fig. 2A** and **Fig. 4B** (lowest density bins excluded). Bins with vL5PNs and fL5PNs are yellow. Bins with vL5PNs only are red. Bins with fL5PNs are blue. **B**. Graphical illustration of the extensive overlap of vL5PNs and fL5PNs in M1 versus the more disjoint organization in S1.

## Discussion

Currently, vM1 is considered to be the major source of cortical influence over vibrissa protraction (6). In turn, vS1 is considered to be the major source of cortical influence over vibrissa retraction (6). Our results suggest two major alterations of this perspective. First, we find that a surprisingly diverse set of cortical areas is the origin of disynaptic control over the MNs that innervate a single protractor muscle. Second, we find that vS1, as well as vM1, are major sources of cortical influence over vibrissa protraction. These new perspectives are discussed in separate sections below.

Before proceeding, we will review some of the technical features of the approach we employed. We injected a single vibrissa muscle with the N2c strain of rabies virus. Retrograde transport of the virus infected ∼30% of the MNs that innervate that muscle. We saw no evidence that rabies virus infected MNs that innervate muscles of other vibrissae, or even the retractor muscles of the C3 vibrissa. Even after extending the survival time to infect 2nd- and 3rd-order neurons, the infected MNs remained confined to those known to innervate the injected protractor muscle **(Fig. S1F)**. Furthermore, the size, shape and location of the MNs infected by rabies virus suggest that our sample is representative of the entire pool of MNs that innervate the C3 protractor muscle (12).

The next important technical feature is that we observed 2nd-order neurons in all of the twenty-one premotor areas of the brainstem **(Fig. S5A-B)** that are known to influence vibrissa protraction (12-19). In addition, we observed no sites of spurious labeling. These observations emphasize that retrograde transsynaptic transport of the N2c strain of rabies virus is both highly specific in terms of infecting only the MNs of the injected protractor muscle, and appropriately robust in its retrograde transsynaptic transport to all excitatory and inhibitory preMNs that innervate the injected muscle.

Three additional findings indicate that this robustness in transsynaptic transport extends to the infection of 3rd-order neurons. First, the 3rd-order neurons in the cerebral cortex that were infected with rabies virus were restricted to cortical layer 5 (vL5PNs), the known source of descending cortical output to subcortical regions. Second, the area of densest cortical labeling in vM1 corresponds to the peak site where stimulation drives vibrissa protraction (6, 18, 22, 23). Third, the peak of the cortical labeling in vS1 aligns with the barrel column corresponding to the injected vibrissa muscle **(Fig. 2E-F)**. In essence, our approach enabled us to generate a comprehensive map of the 3rd-order neurons in the cerebral cortex with disynaptic connections to the MNs of a single vibrissa muscle **(Fig. 6A-B)**.

**Figure 6.**
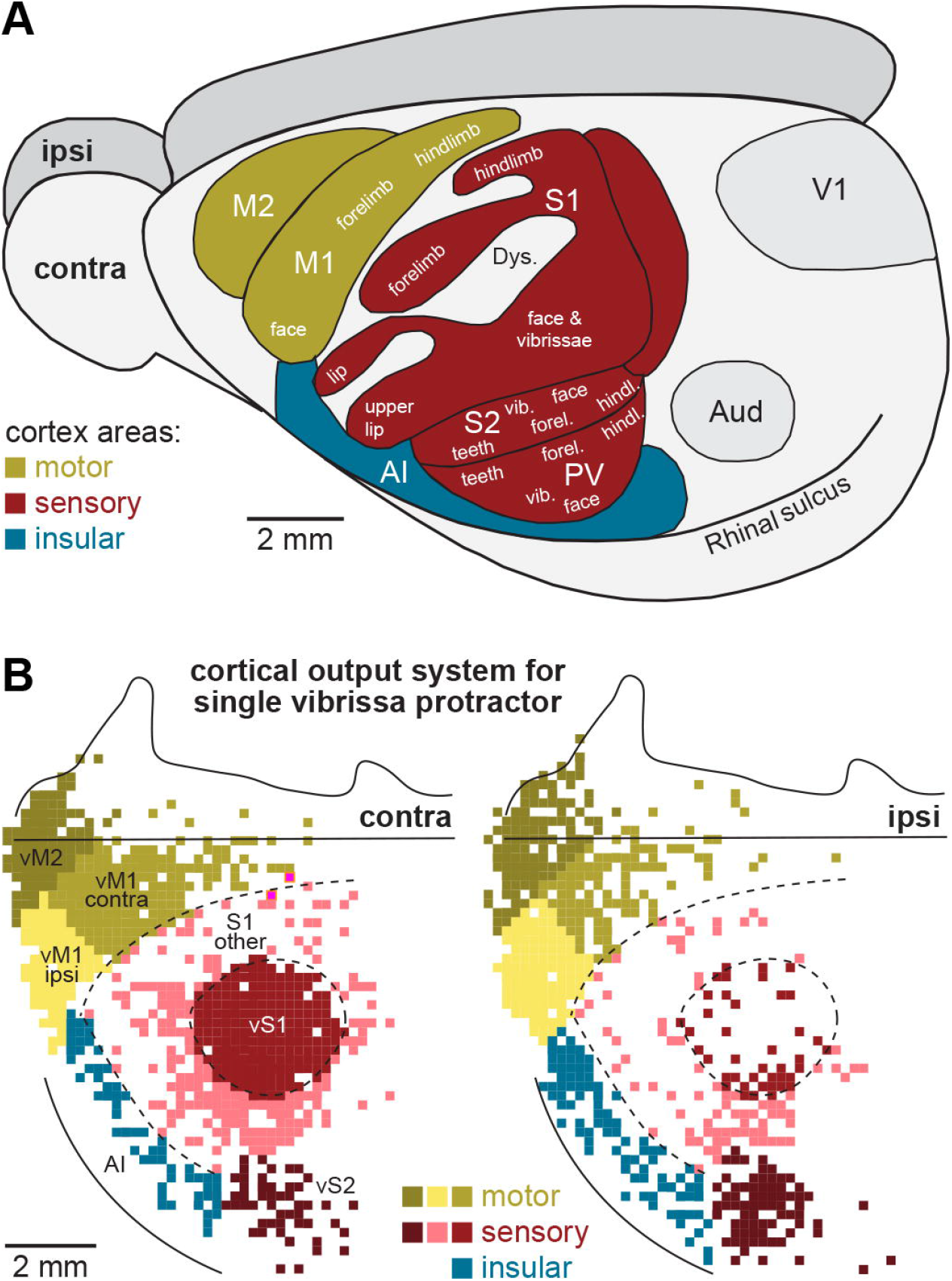
Cortical output system that controls a single vibrissa protractor muscle. **A**. Schematic of cortical areas in the lateral hemisphere of the rat cerebral cortex (modified from (34)). **B**. Regions of the rat cerebral cortex (left: contralateral, right: ipsilateral) with vL5PNs the control protraction of the C3 vibrissa. The functional domains of cortical areas with L5PNs are color-coded.

### Widespread cortical origin of motor commands to vibrissa muscles

It’s long been known that multiple cortical areas are the origin of descending output to the spinal cord and brainstem. Some of these descending cortical output systems are involved in motor processing, whereas others are involved in sensory processing. Our approach enabled us to distinguish between these two general types of efferents, and to identify the cortical neurons that innervate the last-order interneurons that are the major source of input to the MNs of a single vibrissa muscle.

One clear outcome of this demonstration is that more than 2/3rds of the vL5PNs that control a single vibrissa muscle originate from areas outside of M1. These areas are widely dispersed in frontal and parietal cortex. This observation emphasizes the potential importance of descending motor commands from non-primary motor areas. However, the fraction of the disynaptic output from rat M2 is surprisingly small, especially when compared to what is observed in macaques (24). In fact, this is one striking difference between the cortical motor systems in rodents and non-human primates.

More than a third of the vL5PNs originate from somatosensory areas, such as the barrel field (vS1). The barrel field of rodents has been long considered as a prototypic model system for studying sensory processing at the level of the cerebral cortex in mammals (5). Even so, we find that the number of vL5PNs in vS1, and even their peak density, rivals the number and peak density of vL5PNs in vM1. Thus, our results emphasize the importance of the barrel field in the processing of motor output (see also (6) and our discussion below).

Another striking finding of our study is the widespread distribution of vL5PNs within M1. Although vL5PNs are most concentrated in vM1, they are found throughout the entire body map of M1 **(Fig. 6A)**. This is especially well illustrated in our maps **(Fig. 2A)**, which show vL5PNs located as far outside as the hindlimb representation **(Fig. 6B)**. This observation is consistent with prior studies, which found that vibrissae movements can be evoked from stimulation throughout the body map of M1, including the hindlimb representation (25). It is also consistent with electrophysiological data that single L5PNs in rodent M1 can influence the activity of vibrissa muscles and other muscles widely distributed throughout the body (17). We can only speculate on the functional significance of this extreme intermingling. However, it may provide a neural substrate that assists the rodent motor system in developing synergies for coordinated movements of muscles in widely separated regions of the body.

### vM1, vS1 and reciprocal control of vibrissa muscles

It is well known that stimulation of vM1 evokes vibrissa *protraction*, whereas stimulation of vS1 evokes vibrissa *retraction* (22, 23). Similarly, stimulation of the reticular nuclei that receive input from vM1 evokes vibrissa protraction, whereas stimulation of the trigeminal nuclei that receive input from vS1 evoke vibrissa retraction (2, 23). These findings have led to the current model that the vM1→reticular pathway is concerned with the control of vibrissa protraction and that the vS1→trigeminal pathway is concerned with vibrissa retraction **(Fig. 7A)**. How then can one explain our finding that vS1 is a substantial source of vL5PNs with disynaptic connections to the MNs that innervate a protractor muscle?

**Figure 7.**
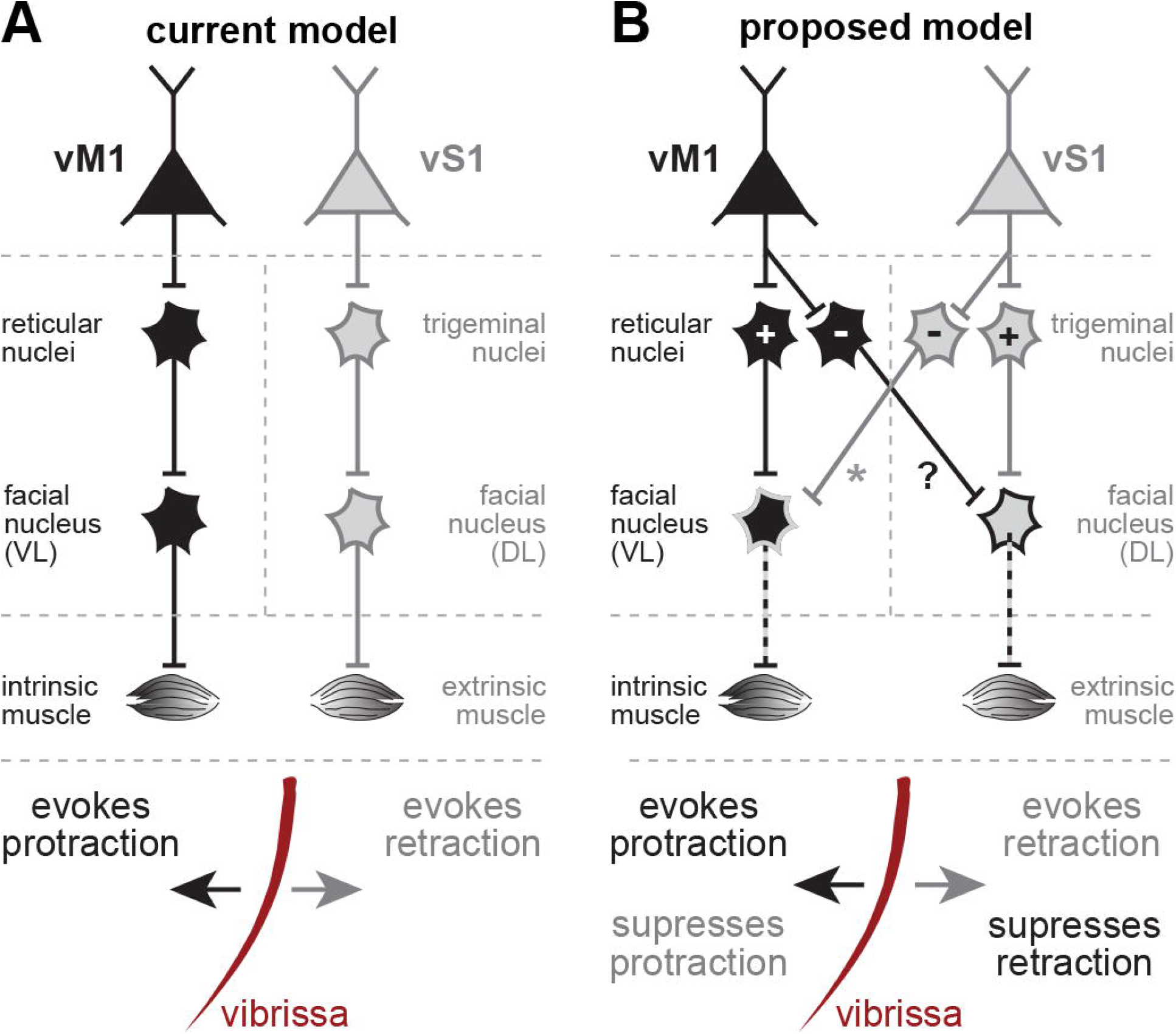
Current and proposed models for cortical control of vibrissa movement. **A**. In the current model (6), vibrissa protraction and retraction are controlled via separate cortical areas and output pathways. The vM1→reticular pathway to the MNs of intrinsic muscles in the ventrolateral (VL) facial nucleus promotes protraction (black), whereas the vS1→trigeminal pathway to the MNs of extrinsic muscles the dorsolateral (DL) facial nucleus promotes retraction (grey). This model leaves unexplained the substantial numbers of vL5PNs labeled in vS1 after injection of the C3 protractor muscle (see main text). **B**. In the proposed model, we add last-order inhibitory interneurons to both the trigeminal and reticular pathways. This model enables the vS1→trigeminal pathway to excite the MNs of retractor muscles and inhibit the MNs of protractor muscles. Similarly, it enables the vM1→reticular pathway to excite the MNs of protractor muscles and inhibit the MNs of retractor muscles. This model explains the substantial labeling in vS1 after injection of the C3 protractor muscle (the asterisks denotes the pathway demonstrated by the current results). The question mark denotes a predicted pathway (see main text).

We believe that this paradox is resolved by a new model **(Fig. 7B)** in which the trigeminal nuclei contain two sets of preMNs, one that excites the MNs of retractor muscles and one that inhibits the MNs of protractor muscles. According to this new model, vL5PNs in vS1that project to the trigeminal nuclei connect to both sets of preMNs. Consequently, the vS1→trigeminal pathway excites the MNs of retractor muscles and inhibits the MNs of protractor muscles. In support of this new model, retrograde transsynaptic transport of rabies virus from the protractor muscle labeled a substantial number of preMNs in the trigeminal nuclei (12) **(Fig. S5A-B)**. Moreover, the model predicts that inactivation of vS1 would remove a source of inhibition to the MNs innervating a protractor muscle. In fact, Matyas et al. showed that inactivation of vS1 leads to enhanced vibrissa protraction from vM1 stimulation (23).

In essence, our model proposes that vM1 is a main source of vibrissa protraction, whereas vS1 is a source of reciprocal inhibition leading to suppression of vibrissa protraction. There is evidence that a similar arrangement exists for the cortical control over vibrissa retraction (23, 26). This would require that vL5PNs in vM1 connect to two sets of preMNs in the reticular nuclei, one that excites the MNs of protractor muscles and one that inhibits the MNs of retractor muscles. Thus, the model predicts that retrograde transsynaptic transport of rabies virus from injections into a *retractor muscle* would label substantial numbers of 3rd-order vL5PNs not only in vS1, but also in vM1.

At this point, we do not know whether any of the single vL5PNs in vM1 branch to excite vibrissa protraction and inhibit vibrissa retraction. Nor do we know whether any of the single vL5PNs in vS1 branch to excite vibrissa retraction and inhibit vibrissa protraction. It is also possible that vS1 contains a small set of vL5PNs that excite vibrissa protraction. All of these issues will need to be explored in future studies using double labeling approaches with virus transport from both protractor and retractor muscles, as well as physiological approaches to assess branching patterns.

Despite this uncertainty, our results suggest that vM1 and vS1 are each the source of activation of a vibrissa muscle agonist and reciprocal inhibition of a vibrissa muscle antagonist. Indeed, our findings emphasize the importance of vS1 as a source of descending inhibition over a protractor muscle. As we noted above, the numbers of vL5PNs in contralateral S1 (26.9% of all vL5PNs, **Table 1**) nearly matches the numbers of vL5PNs in contralateral M1 (29.1%, **Table 1**). Our model predicts that the major effect of the descending signal from vS1 is inhibitory. This cortical command could sculpt vibrissae movements and enhance their individuation through active inhibition (10, 27).

Our results from rabies virus transport from EDC suggest that our model of cortical control extends beyond the vibrissal system. Stimulation of fS1 and fM1 has a push-pull outcome for forelimb movements (25) that is comparable to the effects of vS1 and vM1 stimulation. fS1 stimulation evokes ‘retractive’ movements of the forelimbs, including wrist flexion (25). In contrast, fM1 stimulation evokes ‘forward’ movements of the forelimbs, such as elevation of the forepaw and wrist extension (25). Our results show that both fM1 and fS1 contain fL5PNs with disynaptic connections to the MNs of EDC, a forelimb muscle that extends the wrist and spreads the digits. If our proposed model of vM1-vS1 reciprocal control applies to EDC, then it predicts that fM1 is a source disynaptic excitation of EDC and fS1 is a source of disynaptic inhibition of EDC. Thus, paired initiation and suppression of complementary movements may be a general feature of the descending control signals from the rodent M1 and S1.

## Materials and Methods

### Injections

Experimental procedures were conducted at the University of Pittsburgh, and in accordance with the animal welfare guidelines of the Max Planck Society and the National Institutes of Health. Approval was obtained from the Institutional Animal Care and Use and Biosafety Committees. The handling protocols for rabies virus and animals were previously outlined (20, 28), surpassing the recommendations by the Department of Health and Human Services for Biosafety in Microbiological and Biomedical procedures. For injections into vibrissa muscles, male Wistar rats at postnatal days 28-35 (P28-35) were anesthetized with ketamine-xylazine mixture (70/6 mg/kg, i.p.) and placed in a stereotaxic frame (Helmut Saur Laboratories). The fur around the C3 vibrissa was trimmed, and a small incision was made adjacent to the C3 follicle. The rat’s head was tilted 20° upward in the ventral-dorsal direction to enhance visualization of the mystacial pad on the right side of the snout. The C3 vibrissa was visually identified under a surgical stereoscope (Leica MZ6) and marked at the base using a surgical pen. A steel injection needle connected to a 5 mm glass Hamilton syringe was then inserted into the incision, approximately 1 mm below the surface, with a manual manipulator (Narishige Model BE8). 500-700 nL of rabies virus (N2c strain, 1&10^9^ pfu/mL, provided by M. Schnell, Thomas Jefferson University, Philadelphia) was pressure-injected at the base of the follicle under visual guidance. Following injections, rats were allowed to survive for 3-5 days, during which no observable symptoms were noted. For EDC muscles, we injected 45-50 µl of the N2c strain of rabies virus (>10^8^ pfu/mL) into a single EDC muscle on the right forelimb of Sprague Dawley rats (24). Starting 3 days post injections, rats were transcardially perfused with Phosphate Buffer (PB) at 12 hour intervals. Brains were removed and fixed in 4% paraformaldehyde (PFA) overnight.

### Histology

For vibrissa injections, brains were cut coronally into two parts, a frontal part containing cortex and a caudal part containing cerebellum and brainstem. The caudal part was cut into consecutive 50 µm thick coronal vibratome sections. The frontal part was cut into consecutive 50 µm thick coronal or tangential vibratome sections. For tangential sections, right and left hemisphere were blocked separately, and cut approximately tangential to vS1 (at an angle of 45° with respect to the midline). Sections were permeabilized and blocked in 100 mM PB supplemented with 4% normal goat serum (NGS) (Jackson ImmunoResearch Laboratories) and 0.5% Triton X-100 (TX) (Sigma Aldrich) for 2 hours at room temperature. Primary antibodies, including Rabbit anti-NeuN (diluted 1:500, EMD Millipore #mAB377) and Mouse anti-RABV-P 31G10 (diluted 1:1000, 2mg/ml, Thomas Jefferson University), were incubated in PB containing 1% NGS for 48 hours at 4°C. Secondary antibodies, goat anti-mouse IgG Alexa-488 (1:500) and goat anti-rabbit Alexa 647 (1:500), were then incubated for 2-3 hours at room temperature in PB supplemented with 3% NGS and 0.3% TX. Sections were mounted onto glass slides, embedded in Slowfade Gold (Invitrogen), and covered with a coverslip. For EDC injections, brains were sliced coronally, and sections were sequentially arranged and distributed across ten wells. Every tenth section was stained with Nissl stain. All other sections were processed with avidin-biotin peroxidase (Vectastain, Vector Laboratories) to identify neurons infected with rabies virus using a monoclonal antibody targeting the nucleoprotein of the rabies virus.

### Image acquisition

For vibrissa injections, images were acquired by a confocal laser scanning system (Leica Application Suite Advanced Fluorescence SP5; Leica Microsystems) equipped with a tandem scanning system (Resonance Scanner), spectral detectors with hybrid technology (GaAsP photocathode), and mosaic scanning software (Matrix Screener, provided by Frank Sieckmann, Leica Microsystems). Dual-channel mosaic scans of the entire brain sections were conducted using a 10x glycerol-immersion objective (HC PL APO 10x 0.04 N.A) at a resolution of 0.868 × 0.868 µm per pixel (1.7x digital zoom, 8x line average, 8-kHz scanning speed). For EDC injections, images were acquired by a brightfield microscope system (BX51, Olympus) equipped with mosaic scanning software (Surveyor). Mosaic images of the entire brain sections were obtained using a 10x dry objective at a resolution of 0.926 × 0.926 µm per pixel.

### Data analysis

Images were loaded into Amira software (29) to manually mark neuron somata infected with rabies virus. Outlines of the brain sections, and of different areas in brainstem and cortex were manually delineated using the filament editor in Amira (30). Naming of brain areas was derived from Paxinos Rat Brain Atlas (31). Outlines of all coronal sections from each brain were aligned with respect to the midline and in reference to the sagittal brain from Paxinos Rat Brain Atlas. Tangentially cut brain sections were alignment by blood vessel patterns (32). After alignment, outlines were converted into surfaces and smoothed using ‘Smooth Surface’ in Amira (iterations: 20, lambda: 0.6). The hence generated whole brain reconstructions were aligned along their midlines and digitally converted into flat maps of cortex using ReconWin software (33). The separating borders between motor, somatosensory and insular cortex were determined based on cytoarchitecture as revealed by NeuN labeling. Border between motor and sensory cortex: absence and presence of layer 4. Border between sensory and insular cortex: location above rhinal fissure and reduction in layer thickness. Statistical tests and significance values are provided in the Results section. Data is presented as mean ± standard deviation (STD). Statistical analyses were done with MATLAB, Python, and Prism software. All relevant data are available from the authors.

## Acknowledgments

We thank Dr. M. Schnell (Thomas Jefferson University, Philadelphia, PA) for supplying rabies virus, Dr. A. Wandeler (Animal Diseases Research Institute, Nepean, ON, Canada) for supplying antibody for the rabies virus, M. Page (GreatIslandSoftware.com) for developing software for anatomical reconstructions, M. Engelhardt, Y. Li, A. Wendlandt and K. Kanzler for help with neuroanatomical reconstructions, and H. Park and M. Seetharama for developing early versions of analysis routines. Funding was provided by the Max Planck Institute for Neurobiology of Behavior (MO), European Research Council grants 633428 and 101069192 (MO), Deutsche Forschungsgemeinschaft grants SFB 1089 and SPP 2041 (MO), German Federal Ministry of Education and Research grants 01GQ1002 and 01IS18052 (MO), Neuroscience Network North Rhine-Westphalia grant iBehave (MO), NIH grant P40OD010996 (PS), and the DSF Charitable Foundation grant 1805R01 (PS).

## Author Contributions

MO and PS conceived and designed the study. JG and JAR performed the experiments. AM and FO acquired the data. AM analyzed the data and wrote the first draft of the paper. MO and PS wrote the final version of the paper.

