## Supplemental Figures and Tables for "The Cortical Output System that Controls a Single Vibrissa Muscle"

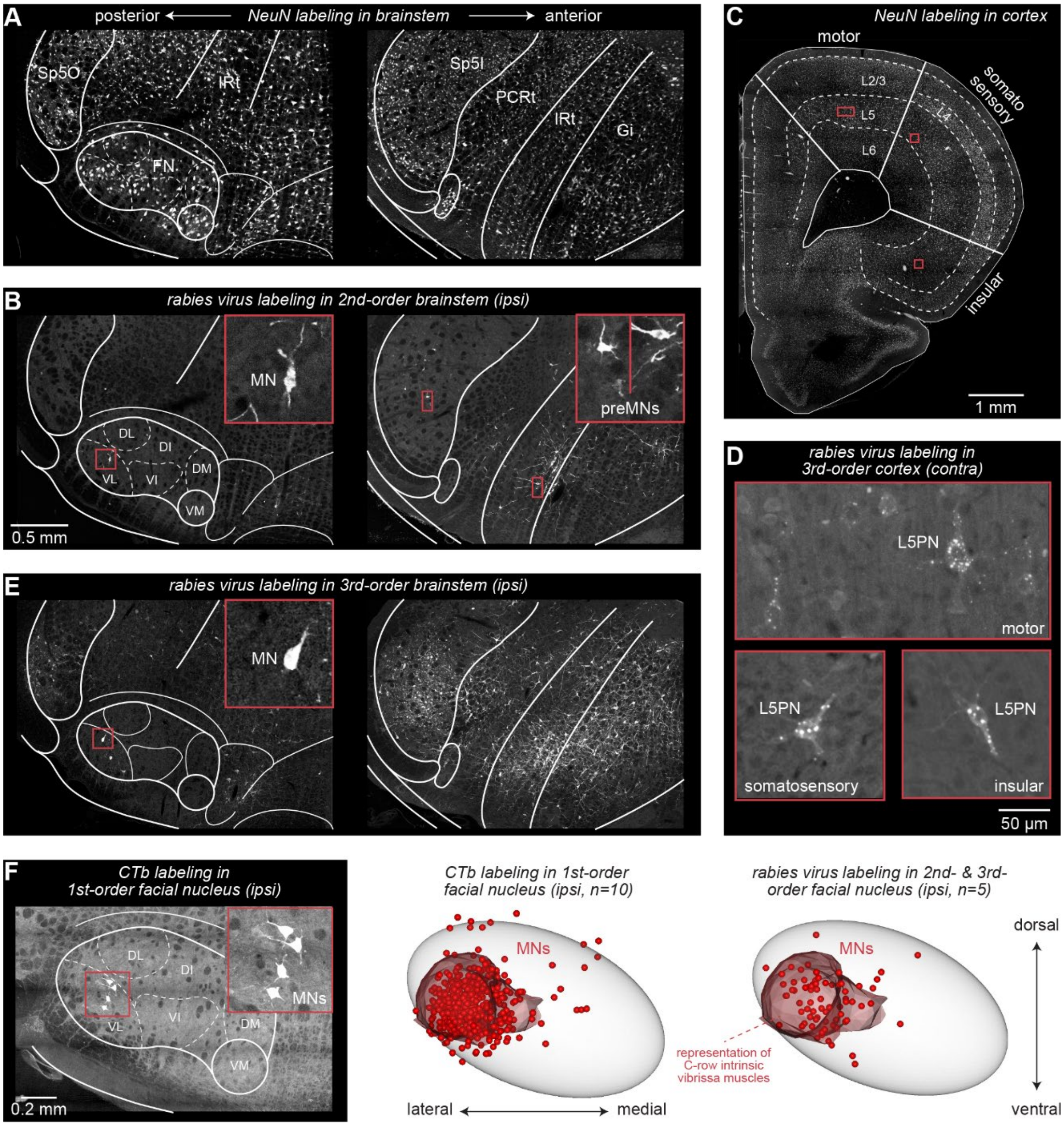


**Fig. S1. Control data for rabies virus transport. A.** Confocal images of 50μm thick coronal section cut through example 2nd-order brainstem. Hemisphere ipsilateral to the injected vibrissa muscles. Lines delineate facial nucleus (FN) and different reticular (PCRt, IRt, Gi) and trigeminal nuclei (Sp5O, Sp5I), identified based on cytoarchitecture via NeuN labeling. **B.** Rabies virus labeling for section from panel A. Dashed lines represent division of the FN into dorsal (D), ventral (V), lateral (L), intermediate (I) and medial (M) parts. Zoom-in to left panel shows MN infected with rabies virus in VL of FN. Zoom-ins to right panel show examples of preMNs infected with rabies virus in trigeminal and reticular nuclei. **C.** Confocal image of coronal section of contralateral cortex from 3rd-order brain. Cytoarchitecture delineates between layers (dashed lines) and cortex areas (solid lines). **D.** Zoom-ins to L5 of motor, somatosensory and insular cortex in panel C show rabies virus infected L5PNs with disynaptic connections to vibrissa MNs. **E.** Same as panel B for example 3rd-order brain. **F.** Left: confocal image of coronal section cut through 1st-order brainstem from rat in which we injected CTb into the C3 vibrissa muscle (1). Zoom-in shows MNs infected by CTb in VL of FN. Center: CTb infected MNs from 10 rats. Right: MNs infected with rabies virus from 5 rats.

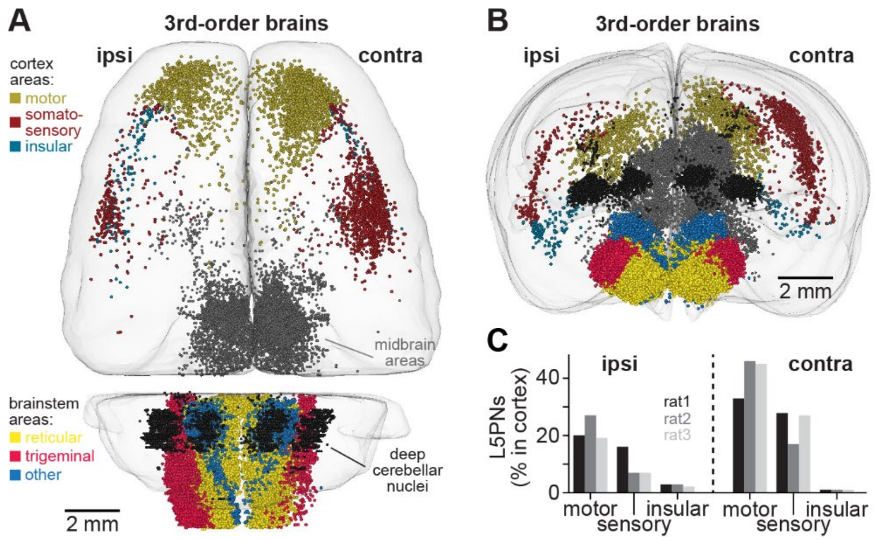


**Fig. S2. Whole-brain reconstructions at 3rd-order spread from vibrissa muscle. A.** Top view onto average brain shows soma locations of all rabies virus infected neurons from three brains where rabies virus did not spread beyond L5. **B.** Coronal view of panel A. **C.** L5PNs in ipsi- vs. contralateral motor, somatosensory and insular cortex across 3rd-order rats (normalized to the total number of L5PNs per rat).

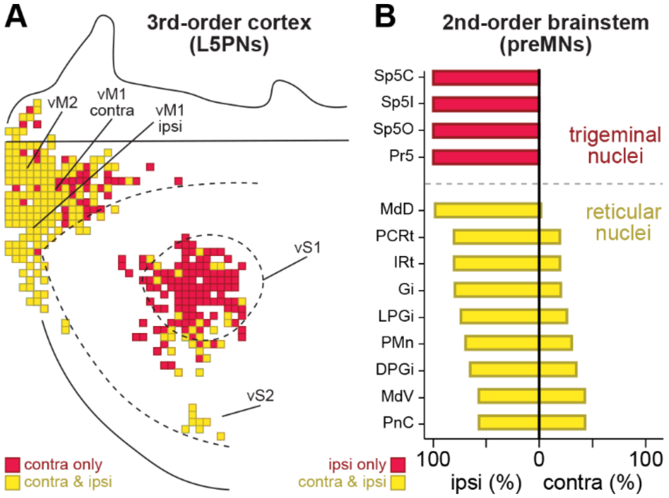


**Fig. S3. Bilateral vs. unilateral control of single intrinsic vibrissa muscle. A.** Overlap between the L5PN density maps from **Fig. 2A-B** (without lowest density bins). Motor areas (vM1/2) and vS2 are bilaterally organized (yellow) – i.e., bins generally comprise L5PNs with disynaptic connections to vibrissa MNs in both hemispheres, whereas vS1 is unilaterally organized (red) – i.e., bins generally comprise L5PNs only contralateral. **B.** Fractions of preMNs in ipsilateral vs. contralateral spinal trigeminal (red) and reticular nuclei (yellow) of 2nd-order brains (n=2) as shown in **Fig. S5**.

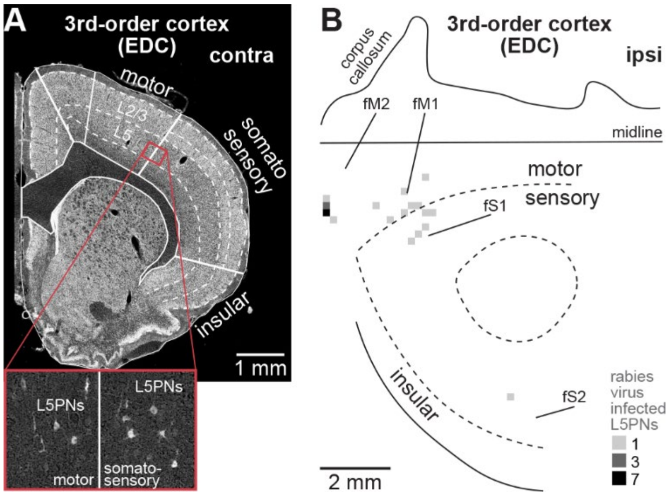


**Fig. S4. 3rd-order spread from single forelimb muscle. A.** Brightfield image of coronal section contralateral to the injected EDC muscle from example 3rd-order brain. We labeled sections for Nissl to delineate between cortical layers (dashed lines) and cortex areas (solid lines). Zoom-in to L5 around motor-to-somatosensory cortex border shows rabies virus infected L5PNs with disynaptic connection to EDC MNs. **B.** Same as Fig. 4B for ipsilateral hemisphere.

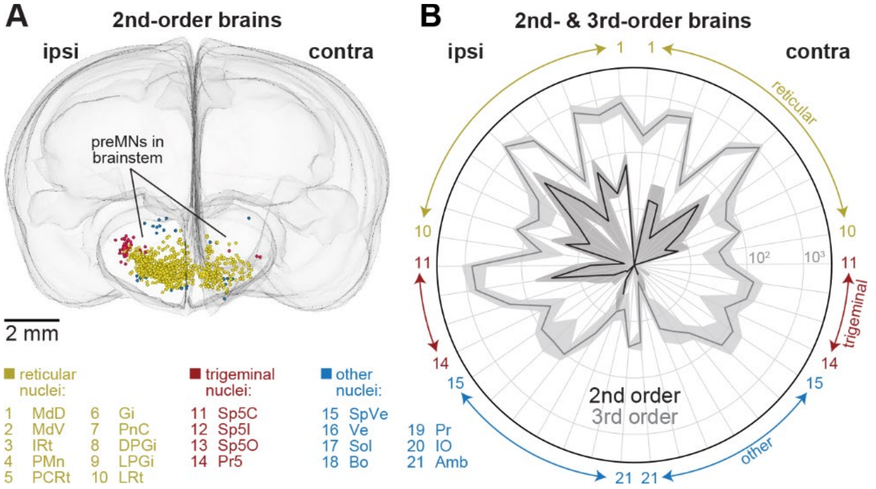


**Fig. S5. Whole-brain reconstructions at 2nd-order spread from vibrissa muscle. A.** Coronal view of average brain shows soma locations of all rabies virus infected neurons from two brains where rabies virus did not spread beyond brainstem. **B.** Radar plots show rabies virus infected preMNs per nuclei in 2nd-order brainstem (n=2, mean ± STD, dark shading), and how the numbers of neurons infected with rabies virus increase in these 21 'premotor' nuclei in 3rd order brainstem (n=3, light shading).

**Table S1:** Neurons infected with rabies virus per area in 3rd-order rat brains (n=3) for injections into C3 intrinsic vibrissa muscle. Non-premotor areas did not comprise neurons infected with rabies virus in 2nd-order brains.

| brainstem | | ipsi | contra |
|---|---|---|---|
| medullary reticular nucleus, dorsal | MdD | 2323 | 595 |
| medullary reticular nucleus, ventral | MdV | 1608 | 1580 |
| intermediate reticular nucleus | IRt | 2404 | 1870 |
| paramedian reticular nucleus | PMn | 156 | 126 |
| parvicellular reticular nucleus | PCRt | 2057 | 1312 |
| gigantocellular reticular nucleus | Gi | 3826 | 3295 |
| pontine reticular nucleus | PnC | 337 | 293 |
| dorsal paragigantocellular nucleus | DPGi | 224 | 191 |
| lateral paragigantocellular nucleus | LPGi | 450 | 273 |
| lateral reticular nucleus | LRt | 138 | 99 |
| spinal trigeminal nucleus, caudal | Sp5C | 2272 | 129 |
| spinal trigeminal nucleus, interpolaris | Sp5I | 2084 | 412 |
| spinal trigeminal nucleus, oralis | Sp5O | 615 | 357 |
| principal sensory trigeminal nucleus | Pr5 | 135 | 178 |
| spinal vestibular nucleus | SpVe | 299 | 383 |
| vestibular nucleus | Ve | 281 | 382 |
| solitary tract | Sol | 309 | 106 |
| Bötzinger complex | Bo | 73 | 69 |
| prepositus nucleus | Pr | 23 | 49 |
| inferior olive | IO | 19 | 6 |
| ambiguus nucleus | Amb | 85 | 71 |
| non-premotor areas | | 3684 | 2439 |
| | **sum** | **23402** | **14215** |

| midbrain | | ipsi |
|---|---|---|
| zona incerta | ZI | 12 |
| superior colliculus | SC | 956 |
| subthalamic nucleus | SubTN | 1 |
| red nucleus | RN | 20 |
| periaquenductal gray area | pGA | 518 |
| substantia nigra | SN | 41 |
| | **sum** | **1548** |

| cortex (L5) | | ipsi |
|---|---|---|
| motor cortex | M1/M2 | 875 |
| somatosensory cortex | S1/S2 | 293 |
| insular cortex | AI | 120 |
| | **sum** | **1288** |

| cerebellum | | ipsi |
|---|---|---|
| deep cerebellar nucleus, medial | | 866 |
| deep cerebellar nucleus, lateral | | 1186 |
| | **sum** | **2052** |

| | total |
|---|---|
| brainstem | 37617 |
| midbrain | 4676 |
| cortex L5 | 4321 |
| cerebrellum | 4502 |
| **sum** | **51116** |

**Table S2:** Same as Table S1, but in percent of total number of neurons infected with rabies virus.

| brainstem | | ipsi | contra |
|---|---|---|---|
| medullary reticular nucleus, dorsal | MdD | 4,54 | 1,16 |
| medullary reticular nucleus, ventral | MdV | 3,15 | 3,09 |
| intermediate reticular nucleus | IRt | 4,70 | 3,66 |
| paramedian reticular nucleus | PMn | 0,31 | 0,25 |
| parvicellular reticular nucleus | PCRt | 4,02 | 2,57 |
| gigantocellular reticular nucleus | Gi | 7,48 | 6,45 |
| pontine reticular nucleus | PnC | 0,66 | 0,57 |
| dorsal paragigantocellular nucleus | DPGi | 0,44 | 0,37 |
| lateral paragigantocellular nucleus | LPGi | 0,88 | 0,53 |
| lateral reticular nucleus | LRt | 0,27 | 0,19 |
| spinal trigeminal nucleus, caudal | Sp5C | 4,44 | 0,25 |
| spinal trigeminal nucleus, interpolaris | Sp5I | 4,08 | 0,81 |
| spinal trigeminal nucleus, oralis | Sp5O | 1,20 | 0,70 |
| principal sensory trigeminal nucleus | Pr5 | 0,26 | 0,35 |
| spinal vestibular nucleus | SpVe | 0,58 | 0,75 |
| vestibular nucleus | Ve | 0,55 | 0,75 |
| solitary tract | Sol | 0,60 | 0,21 |
| Bötzinger complex | Bo | 0,14 | 0,13 |
| prepositus nucleus | Pr | 0,04 | 0,10 |
| inferior olive | IO | 0,04 | 0,01 |
| ambiguus nucleus | Amb | 0,17 | 0,14 |
| non-premotor areas | | 7,21 | 4,77 |
| | **sum** | **45,78** | **27,81** |

| midbrain | | ipsi | contra |
|---|---|---|---|
| zona incerta | ZI | 0,02 | 0,15 |
| superior colliculus | SC | 1,87 | 4,42 |
| subthalamic nucleus | SubTN | 0,00 | 0,00 |
| red nucleus | RN | 0,04 | 0,20 |
| periaquenductal gray area | pGA | 1,01 | 1,14 |
| substantia nigra | SN | 0,08 | 0,21 |
| | **sum** | **3,03** | **6,12** |

| cortex (L5) | | ipsi | contra |
|---|---|---|---|
| motor cortex | M1/M2 | 1,71 | 3,28 |
| somatosensory cortex | S1/S2 | 0,57 | 2,52 |
| insular cortex | AI | 0,23 | 0,14 |
| | **sum** | **2,52** | **5,93** |

| cerebellum | | ipsi | contra |
|---|---|---|---|
| deep cerebellar nucleus, medial | DCNm | 1,69 | 1,14 |
| deep cerebellar nucleus, lateral | DCBl | 2,32 | 3,65 |
| | **sum** | **4,01** | **4,79** |

| | total |
|---|---|
| brainstem | 74 |
| midbrain | 9 |
| cortex L5 | 8 |
| cerebrellum | 9 |
